# Oak regeneration at the arid boundary of the temperate deciduous forest biome: insights from a seeding and watering experiment

**DOI:** 10.1101/2020.08.05.237719

**Authors:** László Erdős, Katalin Szitár, Kinga Öllerer, Gábor Ónodi, Miklós Kertész, Péter Török, Kornél Baráth, Csaba Tölgyesi, Zoltán Bátori, László Somay, Ildikó Orbán, György Kröel-Dulay

## Abstract

Temperate deciduous forests dominated by oaks cover extensive areas in European lowlands. These ecosystems have been under intense anthropogenic use for millennia, thus their natural dynamics, and their regeneration in particular, is still not well understood. Previous studies found that pedunculate oak (*Quercus robur*), one of the most widespread and abundant species in European deciduous forests, regenerates in open habitats and forest edges, but not in closed forest interiors. However, these observations usually come from the core areas of the biome, and much less is known about such processes at its arid boundary, where limiting factors may be different, and climate change may first exert its effects.

In a full factorial field experiment, we tested the effects of different habitats and increased growing season precipitation on the early regeneration of pedunculate oak in a forest-steppe ecosystem in Central Hungary, at the arid boundary of temperate deciduous forests. We planted acorns into three neighbouring habitats (grassland, forest edge, forest interior) and studied seedling emergence and plant performance under ambient weather and additional watering for four years.

In the grassland habitat, seedling emergence was very low, and no seedlings survived by the fourth year. In contrast, seedling emergence was high and similar at forest edges and forest interiors, and was not affected by water addition. Most seedlings survived until the fourth year, with no difference between forest edge and forest interior habitats in numbers, and only minor or transient differences in size (leaf number, height).

The lack of oak regeneration in the grassland contradicts previous reports on successful oak regeneration in open habitats, and may be related to a shift from light limitation to other limiting factors, such as moisture or microclimatic extremes, when moving away from the core of the deciduous forest biome towards its arid boundary. The similar number and performance of seedlings in forest edges and forest interiors may also be related to the decreasing importance of light limitation. The above-average precipitation in the year of seedling emergence (2016) might be a reason why watering had no effect on oak regeneration.

Overall, our results highlight that oak regeneration and thus forest dynamics may be limited by different factors at a biome boundary compared to its core areas. Indeed, this very simple mechanism (inability of oak regeneration in grassland habitats) may contribute to the opening up of the closed forest biome, and the emergence of a biome transition zone.

## 1 Introduction

Temperate deciduous forests characterised by various oak, hornbeam, linden, maple, ash and beech species cover vast areas in Europe (Schultz 2005). They harbour high species richness at the local scale, show high net primary production, and possess considerable carbon sequestration capacity (Pfadenhauer and Klötzli 2014). Though the composition, structure, and abiotic parameters of these forests are well studied (Pfadenhauer and Klötzli 2014), considerable debates regarding their dynamics still exist (e.g. Vera 2002, Svenning 2002, Szabó 2009, Gillian 2016). Uncertainties about the dynamics and especially the natural regeneration of temperate deciduous forests are at least partly due to the fact that most of these forests have been heavily modified by human use during the last couple of millennia, severely compromising natural processes (Ellenberg 1988, Walter and Breckle 1989, Schultz 2005, Kirby and Watkins 2015, Gillian 2016).

Pedunculate oak (*Quercus robur*) is one of the most important tree species in European temperate deciduous forests, dominating lowland forests in a huge belt from Britain to the Ural Mts (Walter and Breckle 1989, Bohn et al. 2004). However, it has been recognised that the natural regeneration of this species is frequently deficient (Shaw 1968 a,b, Reif and Gärtner 2007, Annighöfer et al. 2015). It is well-known that pedunculate oak is a light-demanding species (Annighöfer et al. 2015, Leuschner and Ellenberg 2018). Therefore, its regeneration depends on open or semi-open sites with relatively high light availability, such as forest edges, hedges, openings, and grasslands, and it is not successful in forest interiors (Reif and Gärtner 2007, Leuschner and Ellenberg 2018, reviewed by Bobiec et al. 2018).

Besides light availability, other key factors influencing the regeneration of pedunculate oak include water supply, competition from ground vegetation, zoochory, grazing and browsing (Vander Wall 2001, Annighöfer et al. 2015). Water supply is a critical factor during oak germination and seedling development (Dreyer et al. 1991, Bobiec et al. 2018). While water scarcity is relatively rare in Western Europe and in partially shaded habitats, its effect may be much more important in drier regions and in open habitats (Löf et al. 1998, Reif and Gärtner 2007). Competition heavily influences the survival of seedlings (Jensen and Löf 2017), but it may be reduced in sites where the herb layer is sparse or where ungulates open up the dense sward (Reif and Gärtner 2007). Grazing and browsing can affect seedling survival and performance negatively, but the nutrient reserves of the cotyledon enable oak seedlings to withstand a certain level of defoliation (Frost and Rydin 1997), while thorny shrubs and the high abundandance of other, more palatable species can protect oak seedlings from grazing and browsing (Bakker et al. 2004, Jensen et al. 2012). To sum it up, an area ideal for pedunculate oak regeneration has been described as providing sufficient moisture and consisting of a mosaic of forests, thickets, shrubs, solitary trees, grasslands, and the ecotones between these habitats (Vera 2000, Bakker et al. 2004, Bobiec et al. 2018).

Biome boundaries are regions where several species reach their distributional limits, and there is a major shift in the physiognomy of the vegetation (Walter 1985, Gosz and Sharpe 1989, Neilson 1993, Peters et al. 2006, Pinto-Ledezma et al. 2018). In these transitional zones, patches from both adjoining biomes form a mosaic pattern. Constraints operating in the transitional zones are typically different from those operating within the core areas of the biomes (Gosz and Sharpe 1989, Risser 1995). In addition, species that are dominant in the core area of the biome may become limited to specific habitats (with special microclimates) towards the biome boundary (Gosz 1992, 1993, Neilson 1993). Environmental changes, including climate change, are likely to substantially affect biome boundaries (Gosz and Sharpe 1989, Allen and Breshears 1998, Frelich and Reich 2010). Germination and establishment may be critically affected, resultig in altered dynamic processes in biome boundaries (Gosz 1992, Risser 1995, Erdős et al. 2018).

The forest-steppe zone is at the arid boundary of the temperate deciduous forest biome: as, largely due to climatic constraints, closed-canopy forests open up and gradually give way to grasslands, a mosaic of woody and herbaceous habitats emerges (Wesche et al. 2016, Erdős et al. 2018). Pedunculate oak is a major constituent not only in the deciduous forest biome of Europe, but also in these mosaic ecosystems (Molnár et al. 2012, Erdős et al. 2018). While the regeneration of pedunculate oak has been intensively studied within the core areas of the deciduous forest biome, oak regeneration patterns at the arid boundary of the biome are mostly unstudied (Bobiec et al. 2018).

In this study, our objective was to understand the effects of different habitats (forest interior, forest edge, and grassland) and watering on oak germination and early seedling performance. The experimental area lies at the arid boundary of the deciduous forest biome, where growing season precipitation strongly constrains woody vegetation, therefore we expected that the natural regeneration of oak heavily depends on the amount of precipitation.

Accordingly, our hypothesis was that oak seedling emergence and growth would be positively affected by water addition, especially in grasslands, where evapotranspiration and thus water limitation is highest. Furthermore, in line with previous studies we also hypothesised that seedling emergence and performance would be high in grasslands (only when watered) and in forest edges, but lower and declining through time in forest interiors, because of light limitation.

## 2 Materials and methods

### 2.1 Study area

The Kiskunság Sand Ridge in Central Hungary lies at the arid boundary of the temperate deciduous forest biome. The area is the most arid part of the Carpathian Basin, with a mean annual temperature of 10.5 °C (17.4 °C in the growing season from April to September), and a mean annual precipitation of 530 mm (310 mm in the growing season) (Dövényi 2010). The area is characterised by stabilised calcareous sand dunes, with humus-poor sandy soils (Várallyay 1993). Due to a combination of semiarid climate and coarse-textured sandy soil, forests open up and the potential vegetation is forest-steppe, with both forests and grasslands being natural and permanent elements of the landscape, and forming a mosaic (Erdős et al. 2018).

Pedunculate oak, a characteristic species of the temperate deciduous forests, is also present in this forest-steppe mosaic (Rédei et al. 2020), although its abundance is highly variable and is strongly affected by land use in the past centuries (Biró et al 2013, Erdős et al. 2015). The study area is located near Fülöpháza, Central Hungary; N 46°52’, E 19°25’) (Fig. 1a), where pedunculate oak is currently relatively rare, most likely due to previous land use, but the species is a typical component in several forest-steppe areas in the region.

**Fig. 1.**
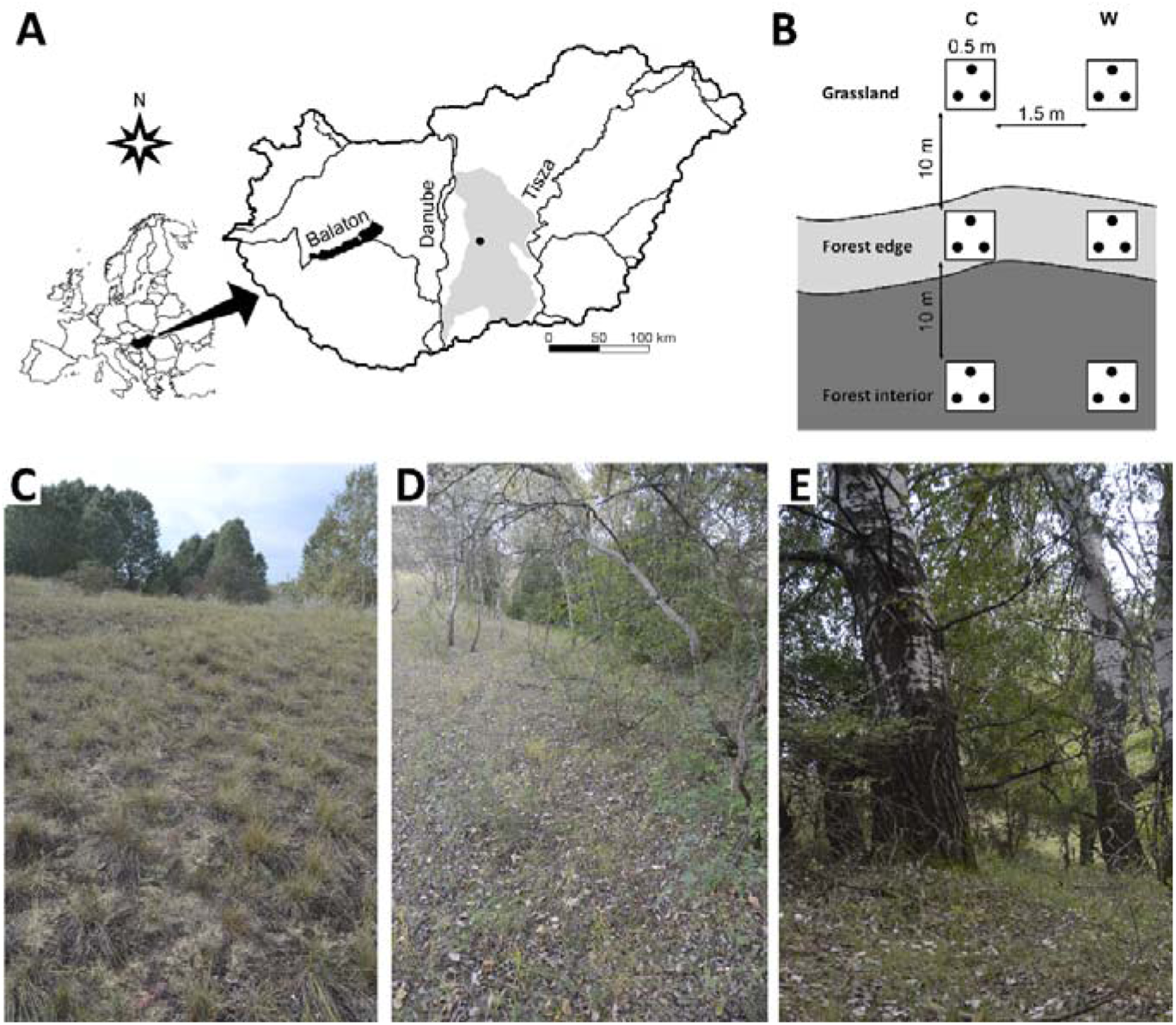
The position of the study area (black dot) in the Kiskunság Sand Ridge (grey shading) (A), the experimental design with oak acorns (black dots) in the 0.5 m × 0.5 m plots in the three habitats under study (C: control plots, W: watered plots) (B), the grassland habitat (C), the forest edge habitat (D), and the forest interior habitat (E).

The forest component of the vegetation mosaic at the study area is represented by the juniper-poplar forest *Junipero-Populetum albae*. The canopy layer is formed mainly by 12-15 m tall *Populus alba* individuals. The shrub layer is dominated by *Juniperus communis* and *Crataegus monogyna*. The most frequent species of the herb layer are *Asparagus officinalis, Carex flacca, C. liparicarpos, Poa angustifolia*, and the seedlings of trees and shrubs.

Among the various grassland communities of the study area, the open perennial sand grassland *Festucetum vaginatae* is the most widespread. Its dominant species are *Festuca vaginata, Stipa borysthenica*, and *S. capillata*, while *Alkanna tinctoria, Dianthus serotinus, Euphorbia seguieriana, Fumana procumbens*, and *Poa bulbosa* are also common.

The contact zones of the forest patches and the grasslands host specific edge communities with various shrubs (e.g., *Berberis vulgaris, Crataegus monogyna, Juniperus communis*) and a high density of *Populus alba* saplings. The most frequent and abundant species of the herb layer include *Calamagrostis epigeios, Festuca rupicola, Pimpinella saxifraga*, and *Taraxacum laevigatum*.

The study area belongs to the Kiskunság National Park; it is strictly protected, and every major human activity except research and controlled tourism has been banned since 1975. The browsing pressure by native ungulates (mostly roe deer) is relatively low. The study area is part of the KISKUN Long-term Ecological Research platform (KISKUN LTER, https://deims.org/124f227a-787d-4378-bc29-aa94f29e1732).

The plant species names follow Király (2009), while the plant community names are used according to Borhidi et al. (2012).

### 2.2 Experimental design

*Quercus robur* acorns were collected in October 2015 from a nearby patch of seed producing oaks. To exclude acorns with reduced viability, we carried out visual inspection and a float test. The float test is reliable in identifying aborted, diseased, insect-infested or otherwise damaged acorns (Gribko and Jones 1995).

Sixteen sites were selected within a ca. 400 m × 1100 m area in a natural forest-grassland mosaic. For each site, three habitats were defined: forest interior (within the forest patch, 10 m from the forest edge), forest edge (the zone outside of the outermost tree trunks but still under the canopy, on the northern side of forest patches), and grassland (a neighbouring treeless area, 10 m from the edge). At each habitat, two 0.5 m × 0.5 m plots were designated in a row parallel to the forest edge. Within both plots, three acorns were planted at a depth of 2 cm in November 2015 (Fig. 1B-E). A total of 288 acorns was used in the experiment (16 sites × 3 habitats × 2 plots × 3 acorns).

At each site and habitat, we applied two precipitation treatments in the two plots: one plot received ambient precipitation (control), while the other plot received additional watering ten times between 5 April and 6 September in their first year (2016). Watering was started in April, because temperature is low until March (ca. 6 °C mean temperature in March) and no water limitation occurs during wintertime. For watering we used rainwater collected nearby, and the amount added corresponded to 15 mm precipitation each time, resulting in a total of 150 mm watering during the year. The additional watering was 36.5% of the natural precipitation in the growing season and 20.2% of the yearly precipitation in 2016.

Seedlings were individually censused every two or three weeks in the first year. The performance of the seedlings was measured near the end of the growing season of the first and the fourth years (19 September 2016 and 25 September 2019, respectively), by registering the following parameters for each plot: (1) the number of living seedlings, (2) the number of leaves per living seedling and (3) the height of the living seedlings.

During the growing season of 2016, we measured the volumetric soil moisture content of the upper 20 cm every two or three weeks from 5 April till 6 September, using FieldScout TDR300 Soil Moisture Meter (Spectrum Technologies Inc). We measured soil water content before watering at each site and five hours after watering in three a priori chosen sites. These two measurements aimed at assessing the longer (ca. two-week-long) and the short-term (right after watering) effects of watering on the soil moisture content. For each 0.5 m × 0.5 m plot, three measurements were done and then averaged. Means for the whole growing season were calculated for each plot.

The Leaf Area Index (LAI) of the woody canopy was estimated above the herbaceous layer (25 cm) using a LAI 2000 Plant Canopy Analyser instrument (LI-COR, Inc., Lincoln, Nebraska). The measurements were conducted in each plot at peak canopy coverage, 30 July 2016, under clear weather conditions. The total cover of the herb layer (percentage of the 0.5 m × 0.5 m plot) was estimated visually on 19 September 2016.

### 2.3 Statistical analyses

All statistical analyses were carried out using the R environment version 3.4.3. (R Core Team 2017). We compared the abiotic conditions of the treated and untreated plots in the three habitat types by using linear mixed-effects (LME) models (nlme package; Pinheiro et al. 2017). We built individual models for soil moisture content before and after watering, LAI, and total herb cover. In the models, habitat type and treatment, and their interaction were used as fixed effects, while site was used as a random effect. As the soil water was measured at only three sites after watering, we analysed the short-term effect of watering by using a linear model where habitat type, watering, and site were all used as fixed effects.

A generalised mixed-effects model (GLMM) with binomial distribution was applied to assess seedling numbers. In these models, the germination success or failure of each acorn was treated as a binary response variable, while habitat type and watering were used as fixed variables, and site as a random variable. Individual models were built for each time. As no seedling survived in the grassland till the fourth year, we did not consider the effect of this habitat type in the respective model.

The effect of habitat type and watering treatment on the leaf number and height of the oak seedlings in both 2016 and 2019 were assessed by applying LME models. In these models, we did not consider the grassland habitat type, as too few seedlings survived in the grassland plots (only 4 individuals by the end of 2016 and none till 2019). Leaf numbers and height data were square-root transformed to meet the homogeneity and normality assumptions of the tests.

We made visual assessments of the residual diagnostic plots to check the assumptions of the tests. For post-hoc pairwise comparisons, we performed Tukey tests using the multcomp package (Hothorn et al. 2016).

## 3 Results

### 3.1 Inherent differences among the studied habitats

The cover of the herb layer was similar in the grassland and the forest edge habitats, while it was much lower in the forest interior habitat (Table 1, Fig. 2a). Note that the cover of the herb layer was relatively low (below 50%) even in the grassland and the forest edge habitats. The LAI of the overstorey vegetation showed marked differences among the habitats, with the lowest value in grasslands, intermediate values at the forest edges, and the highest values in the forest interiors (Fig 2b). Average growing season soil moisture content was the lowest in grasslands, while it was higher and similar at the forest edge and the forest interior habitats (Fig 2c and 2d; control plots).

**Table 1.**
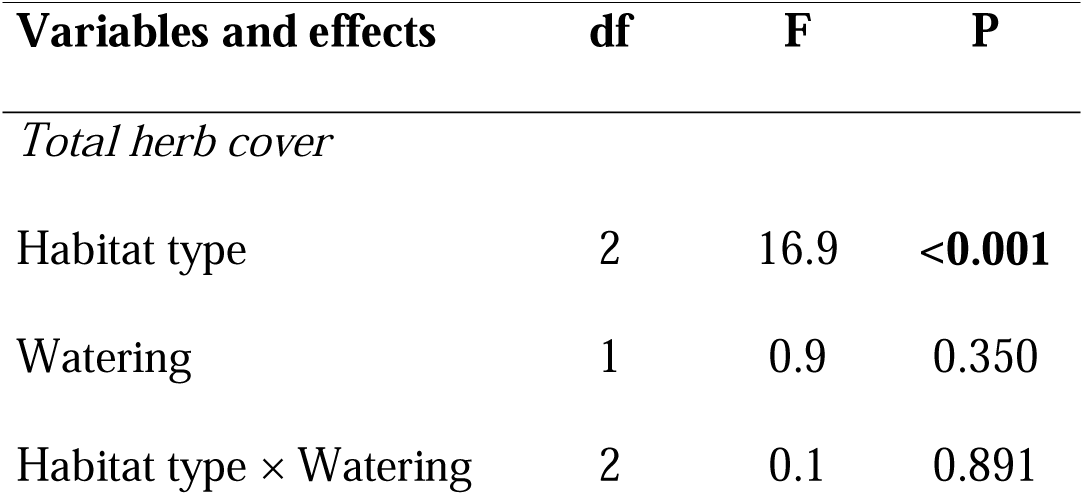
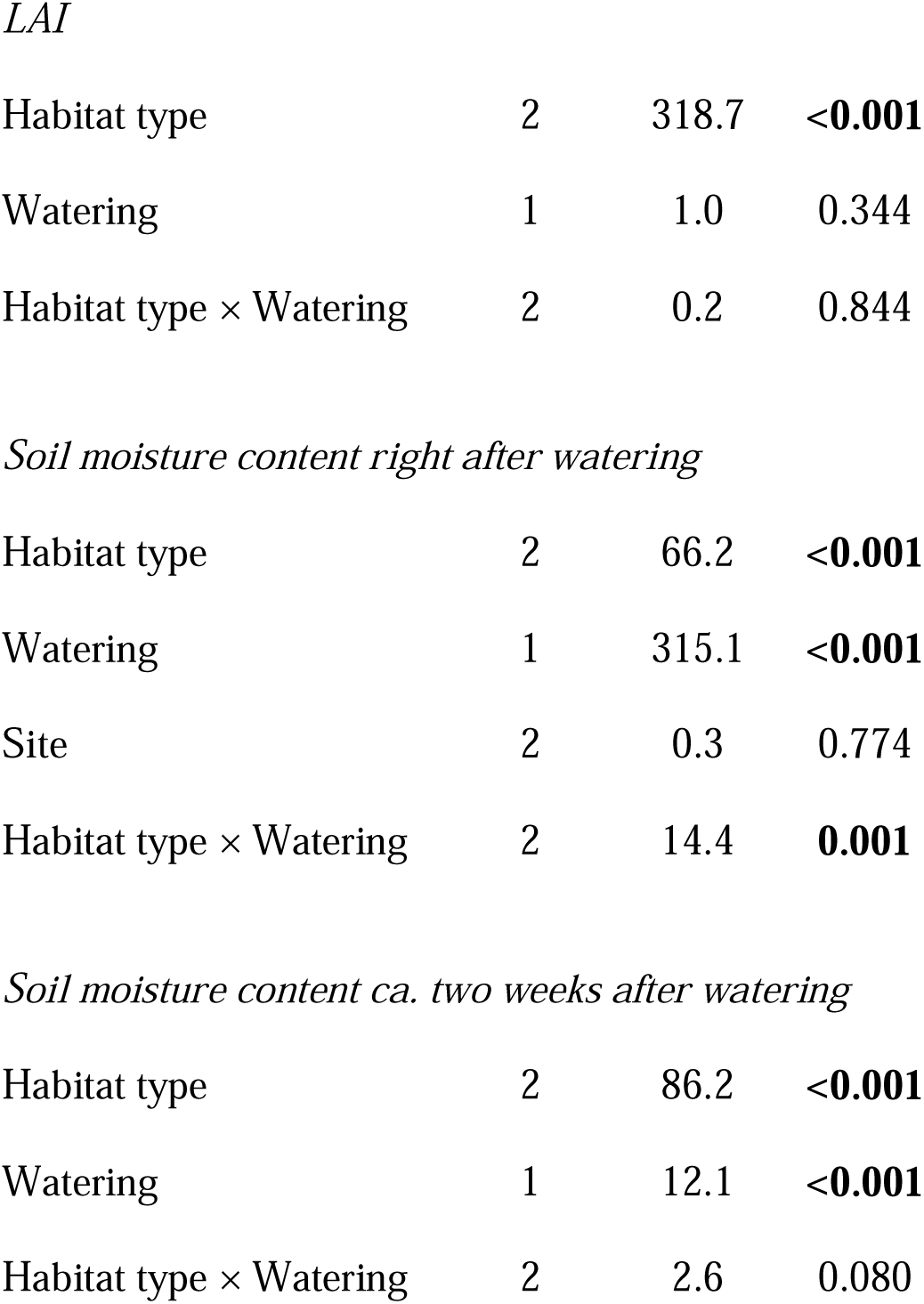
Linear mixed-effects and linear model results of the effects of habitat type, watering on the total cover of the herb layer, soil moisture content before and five hours after watering, and Leaf Area Index (LAI).

| Variables and effects | df | F | P |
| --- | --- | --- | --- |
| *Total herb cover* |  |  |  |
| Habitat type | 2 | 16.9 | <0.001 |
| Watering | 1 | 0.9 | 0.350 |
| Habitat type × Watering | 2 | 0.1 | 0.891 |
| *LAI* |  |  |  |
| Habitat type | 2 | 318.7 | **<0.001** |
| Watering | 1 | 1.0 | 0.344 |
| Habitat type × Watering | 2 | 0.2 | 0.844 |
| *Soil moisture content right after watering* |  |  |  |
| Habitat type | 2 | 66.2 | **<0.001** |
| Watering | 1 | 315.1 | **<0.001** |
| Site | 2 | 0.3 | 0.774 |
| Habitat type × Watering | 2 | 14.4 | **0.001** |
| *Soil moisture content ca. two weeks after watering* |  |  |  |
| Habitat type | 2 | 86.2 | **<0.001** |
| Watering | 1 | 12.1 | **<0.001** |
| Habitat type × Watering | 2 | 2.6 | 0.080 |

**Fig. 2.**
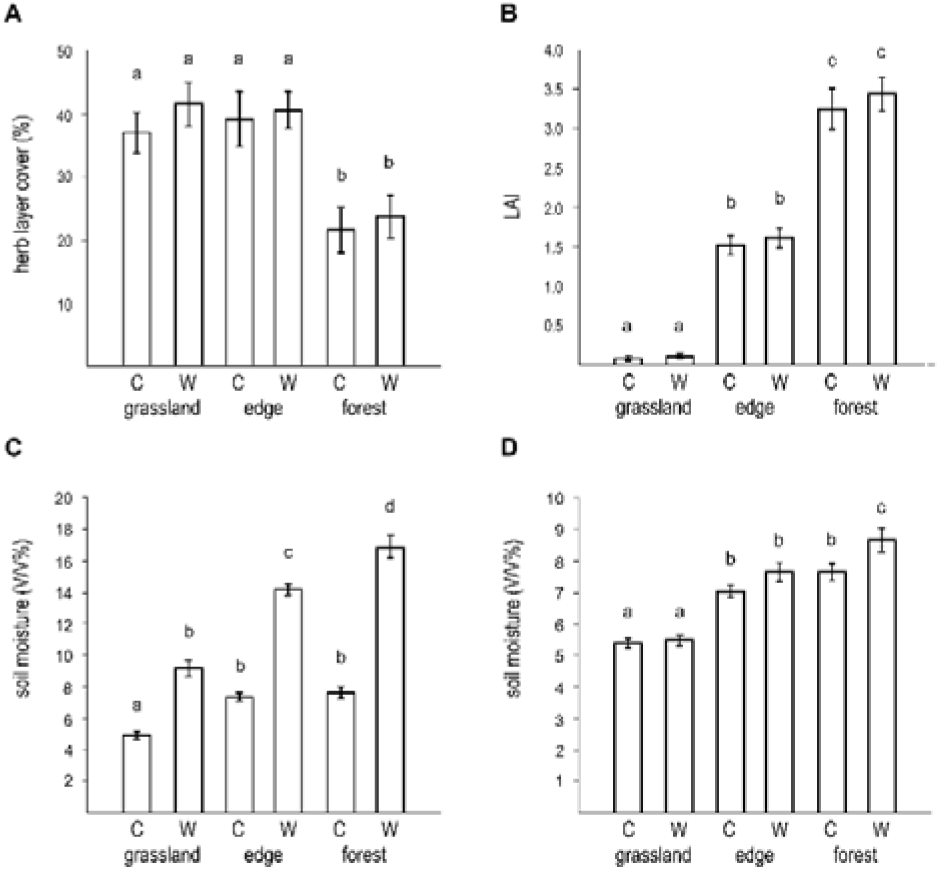
The cover of the herb layer (A), leaf area index (B), soil moisture content five hours after watering (C) and soil moisture content two weeks after watering (D) in the three habitats (grassland, forest edge, and forest interior). C: control plots, W: watered plots.

### 3.2 Effect of watering treatment on soil moisture content

Watering substantially increased soil moisture content in all the three habitats right after watering (Fig 2c), and some of this effect remained even ca. two weeks after watering (before the next watering), although post-hoc tests showed that this was only significant in the forest interior habitats (Fig 2d).

### 3.3 Seedling emergence and survival

Seedling emergence rate was very low in grassland habitats (on average 0.3 acorns germinated out of 3), but was high (on average 2.5 out of 3) and similar in forest edges and forest interiors (Table 2, Fig. 3a). Water addition did not affect the emergence rate (Table 2, Fig 3a). Even the few seedlings that emerged in grasslands died by the fourth year, September 2019 (Fig 3c). Seedling number remained high (around 2) in forest edge and forest interior habitats until September 2019, and was affected neither by habitat (forest edge vs. forest interior) nor by water addition (Table 2, Fig. 3b-c).

**Table 2.**
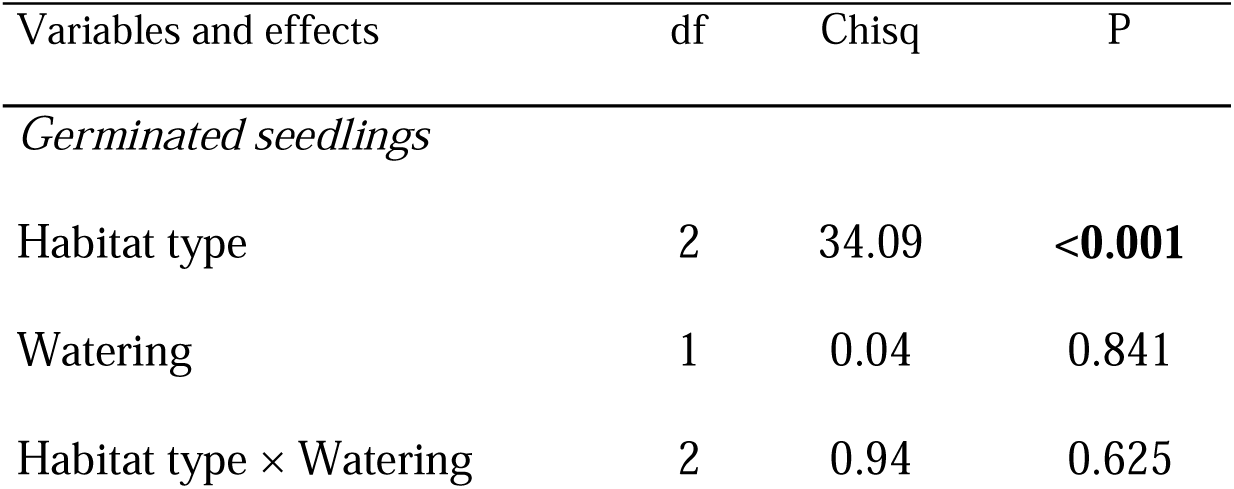
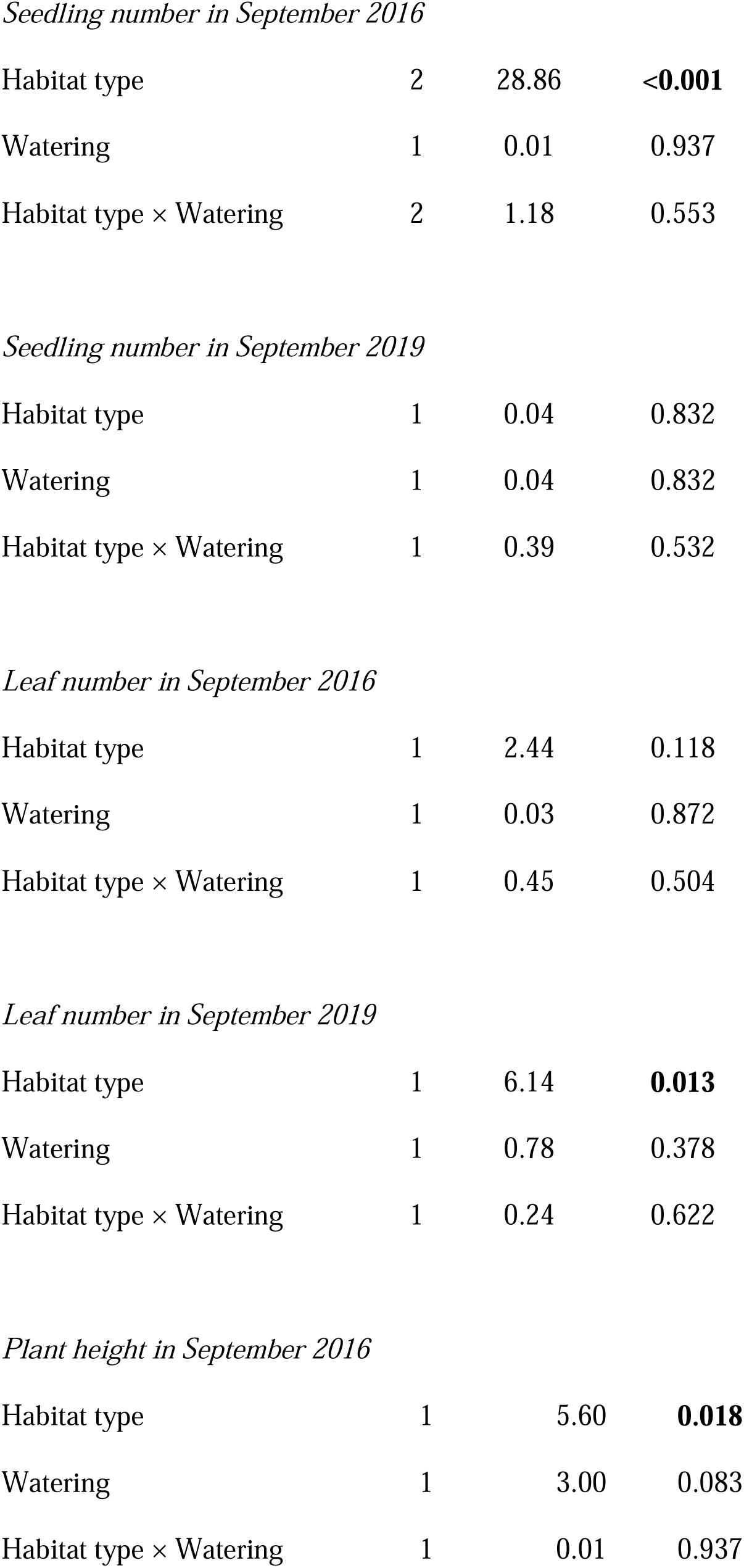
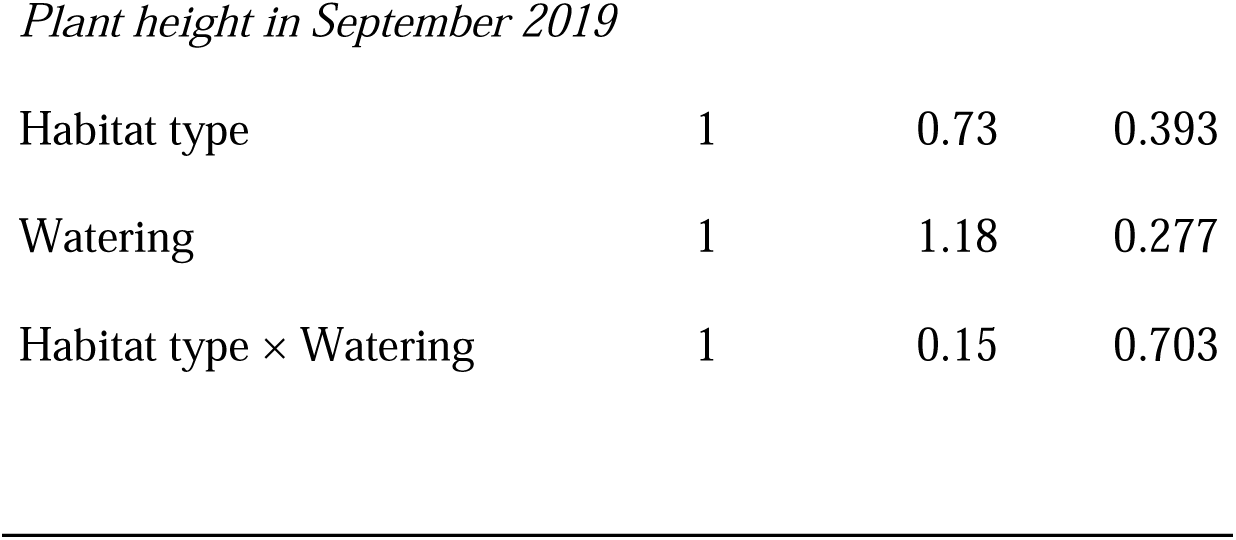
Results of generalised linear mixed-effects model and linear mixed-effects models of habitat type and watering treatment on germinated seedling number, leaf number, and plant height.

| Variables and effects | df | Chisq | P |
| --- | --- | --- | --- |
| *Germinated seedlings* |  |  |  |
| Habitat type | 2 | 34.09 | **<0.001** |
| Watering | 1 | 0.04 | 0.841 |
| Habitat type × Watering | 2 | 0.94 | 0.625 |
| *Seedling number in September 2016* |  |  |  |
| Habitat type | 2 | 28.86 | **<0.001** |
| Watering | 1 | 0.01 | 0.937 |
| Habitat type × Watering | 2 | 1.18 | 0.553 |
| *Seedling number in September 2019* |  |  |  |
| Habitat type | 1 | 0.04 | 0.832 |
| Watering | 1 | 0.04 | 0.832 |
| Habitat type × Watering | 1 | 0.39 | 0.532 |
| *Leaf number in September 2016* |  |  |  |
| Habitat type | 1 | 2.44 | 0.118 |
| Watering | 1 | 0.03 | 0.872 |
| Habitat type × Watering | 1 | 0.45 | 0.504 |
| *Leaf number in September 2019* |  |  |  |
| Habitat type | 1 | 6.14 | **0.013** |
| Watering | 1 | 0.78 | 0.378 |
| Habitat type × Watering | 1 | 0.24 | 0.622 |
| *Plant height in September 2016* |  |  |  |
| Habitat type | 1 | 5.60 | **0.018** |
| Watering | 1 | 3.00 | 0.083 |
| Habitat type × Watering | 1 | 0.01 | 0.937 |
| Plant height in September 2019 |  |  |  |
| Habitat type | 1 | 0.73 | 0.393 |
| Watering | 1 | 1.18 | 0.277 |
| Habitat type × Watering | 1 | 0.15 | 0.703 |

**Fig. 3.**
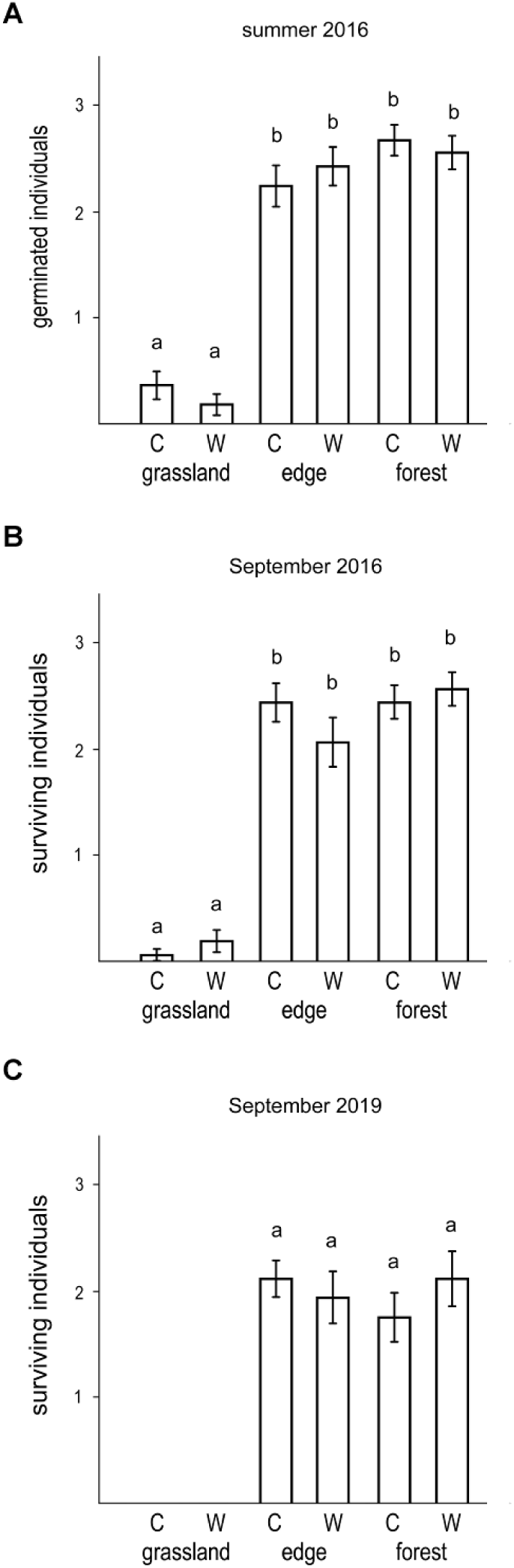
The number of germinated oak individuals (A), individuals that survived until September 2016 (B), and until September 2019 (C) in the three habitats (grassland, forest edge, and forest interior). C: control plots, W: watered plots.

### 3.4 Seedling performance

In September 2016 there was no difference in the leaf number of the seedlings between the forest edge and the forest interior habitats (Table 2, Fig 4a), while in September 2019 seedlings in forest edges had more leaves than seedlings in forest interiors (Table 2, Fig4b). Seedlings were taller in the forest interior habitat in 2016 (Fig. 4c), but there was no difference in plant height between the habitats in 2019 (Fig. 4d). Watering had no effect on leaf number and plant height at either time (Table 2, Fig 4). In general, oak sedlings grew very little from 2016 to 2019, and were still very short and had few leaves at the age of 4 years (Fig 4).

**Fig. 4.**
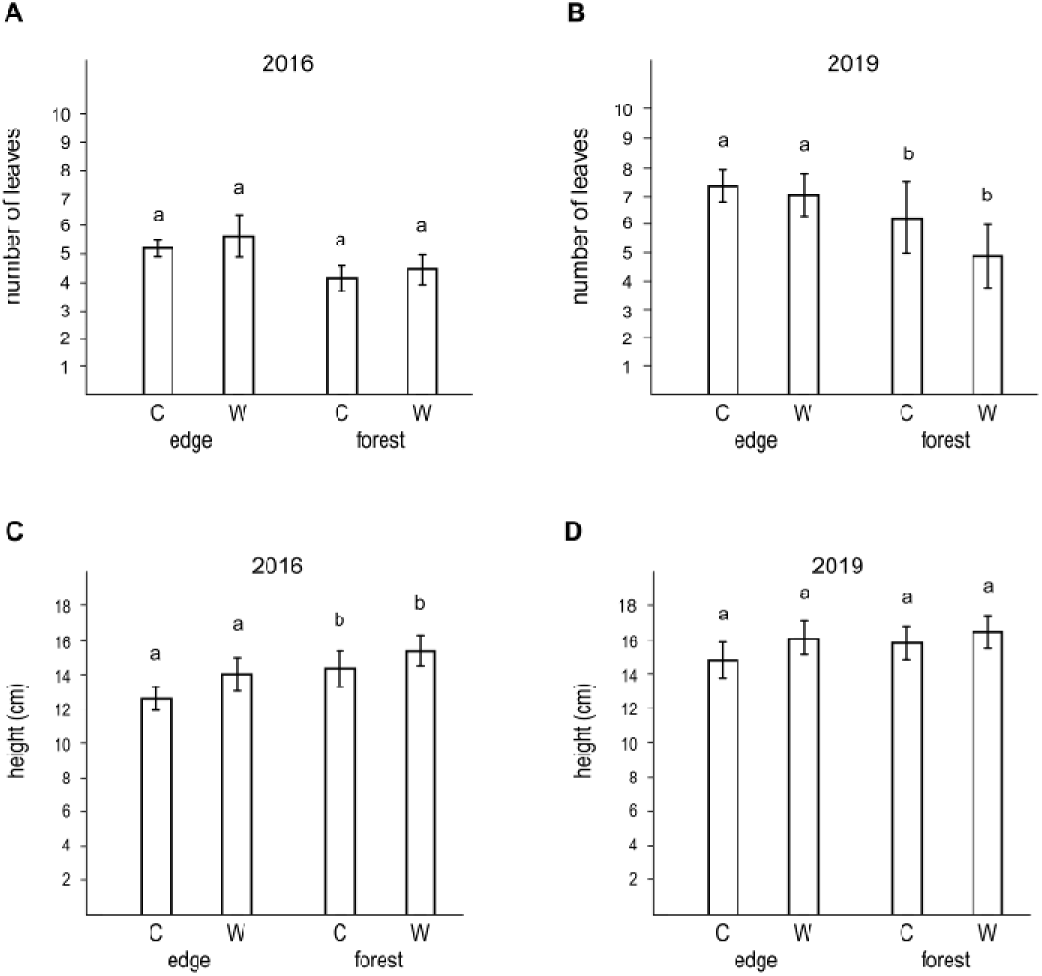
The number of leaves in September 2016 (A), the number of leaves in September 2019 (B), the height of the seedlings in September 2016 (C), and the height of the seedlings in September 2019 (D) in the forest edge and forest interior habitats. C: control plots, W: watered plots.

## 4 Discussion

In contrast to our first hypothesis, watering throughout the growing season did not improve oak seedling emergence and subsequent seedling performance, and this was consistent across all habitat types. Oak seedling emergence and seedling survival were extremely low in the grassland habitat, which contradicts our expectations based on previous reports that pedunculate oak most often regenerates in open or semi-open habitats (Bakker et al. 2004, Bobiec et al. 2018). We did not find a negative effect of the forest interiors compared to forest edges on seedling numbers and performance throughout the four years of the study, while previous studies reported that the shade tolerance of oak seedlings is very low (Lorimer et al. 1994, Welander and Ottosson 1998, Leuschner and Ellenberg 2018). These results suggest that patterns of early oak regeneration at this site at the arid boundary of the temperate deciduous forest biome substantially differ from those previously reported from the core area of the biome. This is most likely related to a shift in oak regeneration from light limitation in the core zone to other limiting factors at the biome boundary.

### 4.1 Effect of watering

Even though we managed to substantially increase soil moisture content during the experiment, excess water had no effect on oak regeneration, which is in striking contrast to our expectation. The pot experiment of van Hees (1997) showed that moist conditions positively affect the height, biomass, and leaf area of *Q. robur* seedlings. The study of Urli et al. (2015) revealed that *Q. robur* seedlings and saplings react sensitively to drought stress in Southwest France and are not able to survive under very dry circumstances. In a Mediterranean mountain environment, Mendoza et al. (2009) found that watering increased the survival of *Q. ilex* and *Q. pyrenaica* seedlings in open and shrubby habitats, while it was not affected under tree canopies, where survival was high even in the absence of watering. In a similar study conducted in Mediterranean ecosystems, Matías et al. (2012a, b) showed that additional watering during the summer is able to increase the survival of *Q. ilex* seedlings in open, shrubby and forest habitats.

The lack of response to watering in our experiment may be related to the fact that 2016 was an unusually wet year. Yearly total precipitation in 2016 was 742 mm, compared to the long-term mean of 530 mm; and growing season precipitation was 410 mm, compared to the long term mean of 310 mm. The fact that even a year of above-average precipitation combined with excess water resulted in very low emergence and no survival in grassland patches suggests that grasslands are truly incapable of supporting oak regeneration in this ecoystem.

In our experiment, watering lasted throughout the growing season, from early April to September. Although we did not assess potential effect of water limitation outside the growing season, the cool temperature combined with usually substantial water in this period (an average 30-50 mm per month, Kovács-Láng et al 2000) makes water limitation unlikely.

### 4.2 Effect of habitat type

In contrast to our hypothesis that the forest edge would represent the best habitat for seedlings while the grassland (due to drought) and the forest interior (due to shade) habitats would be less suitable, we found that seedling emergence and performance were extremely poor in grasslands while they were high in forest edges and forest interiors. Thus, forest edges and forest interiors proved to be similarly suitable for early oak regeneration, despite the strong differences regarding abiotic parameters in these two habitats.

Oak regeneration was absent in the grassland habitat: seedling emergence was extremely low and the few seedlings that did emerge died by September 2019. This finding contradicts earlier studies from the temperate deciduous forest biome. For example, Bakker et al. (2004) found that the survival and performance of pedunculate oak seedlings was better in grasslands than in forest interiors in riverine floodplains of western Europe (Germany, the Netherlands, and Great Britain). Similarly, *Q. robur* is able to colonise abandoned ploughlands and pastures as shown in France (Onaindia et al. 2001) and Poland (Bobiec et al. 2011b). The study of Olrik et al. (2012) showed successful colonisation by pedunculate oak in a heathland in Denmark, while oak can occupy abandoned pastures in Poland and Ukraine (Ziobro et al. 2016). In Belgium, several non-woody vegetation types such as grasslands, ruderal fields and bramble thickets proved to be appropriate for *Q. robur* emergence (Van Uytvanck et al. 2008). Thus it seems that *Q. robur* can easily regenerate in open (i.e., non-woody) habitats in the core areas of the temperate deciduous forest biome (Bobiec et al. 2018).

However, studies from Mediterranean habitats with oak species other than pedunculate oak indicated that oak regeneration may be limited in open habitats. For example, Mendoza et al. (2009) reported from southern Spain that the seedling survival of two Mediterranean oak species, *Q. ilex* and *Q. pyrenaica*, was the lowest in open habitats, while it was much higher under shrubs and in woodlands. Matías et al. (2012b) found that the emergence of *Q. ilex* was very good in open habitats, but the survival of the seedlings was poor in the same habitat, presumably due to drought stress. Similarly, in southern France, Rousset and Lepart (2000) showed that the germination and survival of *Q. humilis* was better under shrubs than in the neighbouring grassland, as shrubs protected the seedlings from drought. In Mediterranean California, the seedling transplantation study of López-Sánchez et al. (2019) revealed that almost all seedlings of *Q. lobata* and *Q. agrifolia* died in the open grassland, while they had significantly higher survival rates under trees and shrubs, where they were more protected from drought stress.

Desiccation is a critical factor during oak germination and seedling growth (Bobiec et al. 2018). Low soil moisture seems to be the most likely cause of poor seedling emergence and performance in our study, besides other factors, discussed below. Water limitation is the most prominent ecological constraint in the centre of the Carpathian Basin, with a semi-arid period during the summer months according to the long-term climate records (Borhidi 1993, Kun 2001). In addition, the sandy soils of the study site have very poor water retention capacity (Várallyay 1993), further decreasing water availability. While water limitation is relatively rare in the western and northern parts of Europe (within the core area of the temperate deciduous forest biome) (Reif and Gärtner 2007), it seems to be of primary importance at the arid boundary of the biome.

Competition with ground vegetation is usually considered one of the most important factors limiting oak regeneration (e.g., Vander Wall 2001, Reif and Gärtner 2007, Annighöfer et al. 2015). However, we think it cannot explain the strikingly poor oak regeneration in grasslands. First, the total cover of the herb layer was very low (40% or even less) in the grasslands of the study. Thus there was ample space for oak seedlings to establish. Second, the cover of the herb layer was similar in the grasslands and the forest edges, yet forest edges had much higher seedling emergence and performance rates.

The poor oak germination and performance of the grassland habitat cannot be explained by browsing or predation either (Bobiec et al. 2018). Browsing pressure is generally low in the area, and we did not see signs of heavy browsing pressure on the seedlings during our regular surveys. Seed predation is also unlikely to differ substantially among the three habitats, due to the small distances (few metres) between the forest interior, forest edge, and grassland plots.

Further factors potentially limiting oak regeneration include high solar radiation and the lack of a humus layer (Nilsson et al. 1996), both of which might have affected seedling emergence and survival in our study. The influence of high solar radiation may be amplified by the very sparse herb layer, and may contribute to the drying of the soil. Regarding the humus layer, the sandy soil of the grassland habitat in the study site is extremely poor in humus: the humus content of the upper 10 cm soil layer can be as low as 0.6%, while it is considerably higher in the forest patches (Bodrogközy 1982, Várallyay 1993, Kröel-Dulay et al. 2019, Tölgyesi et al. 2020).

Microclimatic extremes may also contribute to the poor oak emergence and survival in the grassland habitat. High air temperatures measured near the soil surface in the grassland habitat during summer days (compared to the much cooler forest edges and forest interiors) (e.g., Erdős et al. 2014, Tölgyesi et al. 2020) may damage the tissues and physiological processes of pedunculate oak (Cuza 2018), thus preventing oak regeneration in this habitat.

Our study revealed similarly high early oak regeneration in forest edges and forest interiors. Good oak regeneration within the forest edge habitat fits our hypothesis and is in line with earlier observations regarding habitats optimal for oak regeneration (e.g., Vera 2000, Reif and Gärtner 2007, Bobiec et al. 2018). For example, Herlin and Fry (2000) showed that *Q. robur* is able to establish in forest edges and hedgerows in southern Sweden. Similarly, Bakker et al. (2004) found that edges are optimal habitats for *Q. robur* regeneration throughout northwestern Europe.

We found only small and transient differences between forest edge and forest interior habitats. Seedlings in the forest interiors had fewer leaves than seedlings in forest edges, although the difference was significant only in 2019. This result is in line with earlier studies reporting reduced leaf number in seedlings under shady conditions (e.g., Ziegenhagen and Kausch 1995, Welander and Ottosson 1998). Seedlings were higher in forest interiors than in forest edges in 2016, while no significant difference was found in 2019. Seedlings are usually higher in shady than in sunny habitats (e.g., Ziegenhagen and Kausch 1995, Nilsson et al. 1996, van Hees 1997, Ammer 2003).

The overall similarity of forest interiors and forest edges is surprising given the reported high light requirements of pedunculate oak seedlings. According to Leuschner and Ellenberg (2018), the shade tolerance of *Q. robur* seedlings is very low. Indeed, the regeneration of pedunculate oak depends primarily on non-forest habitats (Bakker et al. 2004, Bobiec et al. 2018). However, it has also been shown that seedlings do tolerate shady conditions during the first few years; that is, their light demand starts to increase only after those initial years (e.g., Welander and Ottosson 1998, Vander Wall 2001, Annighöfer et al. 2015). Bobiec et al. (2011a) found that young oak individuals are increasingly dependent on clearings as they grow up. Ziegenhagen and Kausch (1995) argued that the starch reserves of the young seedlings enable them to survive in shade for a couple of years. Although a negative effect of shading in the forest interiors may easily be seen in the future, the lack of such difference in the first four years is interesting given the above reports on low shade tolerance of pedunculate oak. One possible explanation may be that forest interiors at our site are not as closed as forests in the biome interior (see picture in Fig 1E). Indeed, the LAI of 3-3.5 measured at our forest interiors is lower than that reported for several temperate oak forests in Europe (e.g., Bréda and Granier 1996, Le Dantec et al. 2000, Soudani et al. 2006, Thimonier et al. 2010). Another explanation for the similar performance of oak seedlings at forest edges and forest interiors is that a factor other than light limits growth. A major candidate in these ecosystems can be soil moisture (Várallyay 1993), which may also explain the extremely small size of the 4-year old oak seedlings (14-16 cm).

### 4.3 Differences in oak regeneration between the core area and the arid boundary of the biome

Towards the arid boundary of the temperate deciduous forest biome, the competitive vigour of the woody life-forms decreases (Walter and Breckle 1989, Erdős et al. 2018). As a consequence, forests gradually open up, enabling the emergence of the forest-steppe zone with alternating forest and grassland patches. The poor performance of our seedlings, especially regarding their height, also indicates that conditions are suboptimal for oak regeneration at our site. In our study, oak seedlings were very small after four growing seasons, reaching a mean height of ca. 14-16 cm, which is extremely short compared to seedlings growing under more optimal conditions. Seedling height has been reported to reach 13-20 cm after one (Giertych and Suszka 2010, Devetaković et al. 2019), and 30-60 cm after only two growing seasons (Ammer 2003, Cabral and O’Reilly 2008, Andersen 2010).

## 5 Conclusions

Our study suggests that oak regeneration pattern in this transitional zone differs markedly from what has been described in the core areas of the temperate deciduous forest biome. When one moves from the core areas of the deciduous forest biome towards the arid boundary of the biome, there seems to be a shift from light limitation to other limiting factors, which prevent oak regeneration in grassland patches and restrict it to forest edges, and, potentially, to forest interiors.

In conclusion, our results emphasise that oak regeneration and thus forest dynamics may be limited by different factors at a biome boundary compared to the biome core. Indeed, the lack of tree regeneration in grassland patches may contribute to the opening up of the closed forest biome, and the emergence of the forest-steppe zone.

## Acknowledgements

Funding: This work was supported by the Hungarian Scientific Research Fund (grant number OTKA PD 116114 for László Erdős), the National Research, Development and Innovation Office (grant numbers NKFIH K 119225 and NKFIH KH 129483 for Péter Török, NKFIH K 124796 for Zoltán Bátori, NKFIH PD 132131 for Csaba Tölgyesi, NKFIH K 129068 for György Kröel-Dulay). Kinga Öllerer received further support from the Romanian Academy (grant number RO1567 IBB03/2019). The authors are thankful for the support of the ‘Momentum’ Program of the Hungarian Academy of Sciences.

